# Data-driven burst shape analysis for functional phenotyping of neuronal cultures

**DOI:** 10.1101/2025.09.29.679256

**Authors:** Tim J. Schäfer, Emmanouil Giannakakis, Paul Schmidt-Barbo, Anna Levina, Oleg Vinogradov

## Abstract

Cultures of neurons *in vitro* are instrumental for studying network dynamics in normal and pathological conditions. Mature networks typically exhibit network bursting activity, which has traditionally been quantified by simplified features such as inter-burst intervals and burst durations. While these features advanced the understanding of development, disease phenotypes, and drug effects, they overlook the temporal structure of activity within bursts. Here, we developed a comprehensive framework to quantify burst shapes, the time course of network firing during bursts. Applying this approach to four datasets, including rodent- and human pluripotent stem cell-derived cultures, we show that burst shapes contain rich information about the underlying network dynamics. We quantify this information by using traditional and shape features to classify the recording conditions (types of genetic disorder, presence of pharmacological agents) and demonstrate that shapes significantly increase classification accuracy. We provide a pipeline for burst shape characterization, including simplified features that capture most of the shape information, establishing burst shape as a robust and biologically meaningful marker for functional phenotyping in disease modeling and drug screening.

## Introduction

Synchronized network bursts are a fundamental activity mode of developing neural networks *in vivo* and *in vitro*, arising from the interplay of excitability, connectivity, and dynamic neuron characteristics [1**?** –5]. Neural cell cultures often develop such bursting activity, typically characterized by synchronous firing lasting from tens of milliseconds to a few seconds, followed by long, irregular quiescent intervals. Such activity patterns are observed in cortical, hippocampal, striatal, and spinal cord cultures *in vitro* as well as cultures of both embryonic and induced human pluripotent stem cell (hPSC)-derived neurons [4, 6–9]. Deviation of bursting dynamics statistics from healthy controls *in vitro* is a strong indicator of neuro-developmental disorders [10–13].

Traditionally, population activity in neuronal cultures has been characterized by summary statistics of bursting episodes, such as duration, inter-burst interval, and average firing rate [7, 14, 15]. With the ever-growing availability of tools to monitor network activity [16, 17], these measures have become a common instrument to assess the effects of various mutations, as well as chronic and acute perturbations on individual neurons and networks [8–10, 12, 18–21].

However, the burst shape, the detailed time course of firing rate within the burst, has received comparatively little attention, even though theory, simulations, and experiments all point to a close link between network properties and intra-burst dynamics [6, 11, 22, 23]. Previous works attempted parametric descriptions of burst shapes tailored to individual experiments, including features such as rise, peak, and decay time[9, 15, 24, 25]. While informative, these simple metrics are limited in two key aspects. First, they oversimplify complex activity patterns, such as fluctuating firing rates, rebound firing, and oscillations. Second, summarizing burst shapes into a few numbers per recording conceals the variability of burst shapes within each culture, potentially obscuring biologically meaningful heterogeneity.

To overcome these limitations, we introduce a data-driven framework for normalizing and comparing burst shapes in a standardized way. Applied to four published datasets of rodent- and hPSC-derived neural cultures [9, 15, 25, 26], this framework shows that burst shape information significantly improves the classification of experimental conditions, revealing a close link between intra-burst dynamics and underlying network properties. Our approach enables visualization of burst shapes and extraction of compact, interpretable descriptors that approach the performance of the full data-driven representation. Additionally, we find that genetically similar cultures exhibit similar burst shape patterns, establishing the burst shape as a biologically meaningful marker of network phenotype.

## Results

We aim to maximize the information extracted from multi-electrode array (MEA) recordings obtained under diverse experimental conditions, including pharmacological manipulations and disease-related mutations (Figure 1a). Traditionally, population activity has been described by the set of simple statistical features such as burst duration, inter-burst interval, and firing rates (Figure 1b). Because these measures overlook the temporal structure within bursts, we propose to augment them with burst shape features. In the following, we describe how burst shapes can be represented and compared, and how these representations can be incorporated into statistical models trained to infer experimental conditions. Within this framework, the improvement in classification accuracy quantifies the additional information carried by burst shapes (Figure 1c).

**Fig. 1.**
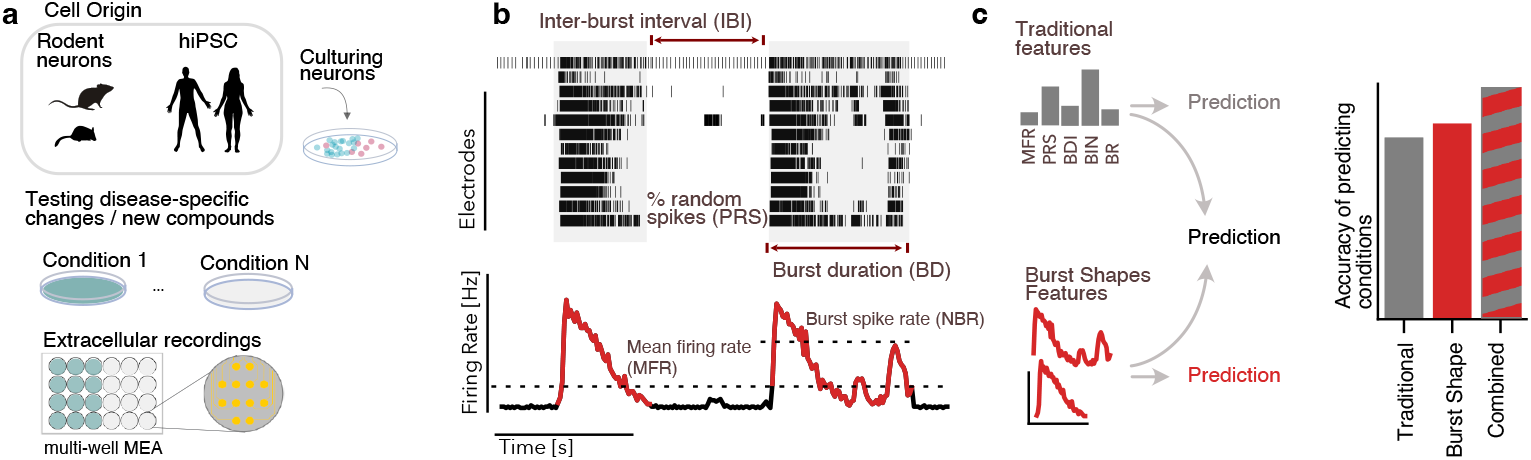
Schematic of evaluating burst shapes as complementary to traditional features for network phenotyping. **(a)** Neural cell cultures, including rodent- and hPSC-derived cells, plated and recorded with multi-electrode arrays (MEAs). **(b)** Traditional descriptors of bursting activity (inter-burst intervals, burst duration, percentage of random spikes, firing rate overall and during bursts) are complemented by burst shape features. **(c)** The contribution of traditional and burst shape features is assessed using models that infer the experimental condition from the different feature subsets.

### Framework for data-driven quantification of burst shapes

We analyzed four datasets of MEA-recorded network activity from neural cultures under various experimental conditions (dataset abbreviations in brackets, see Methods for details): Mouse cortical neurons with blocked inhibition (Inhib. block) [26], hPSC-derived neurons with *CACNA1A* haploinsufficiency modification (*CACNA1A*) [25], hPSC-derived neurons from Kleefstra syndrome patients and controls (Kleefstra) [9], and rat cortical neurons in genetic batches (Rat Cortex) [15]. We detected 31,949 network bursts in 970 recordings and 604 hours of activity (Supplementary Table S1). To ensure reliable burst detection at this scale, we developed an interactive tool for detailed review and parameter tuning (Supplementary Figure S1, detection parameters summarized in Supplementary Table S2).

To compare burst shapes across a wide range of durations (tens of milliseconds to several seconds) and amplitudes (10-3000 Hz), we standardized all burst shapes to unit duration and unit integral (Figure 2a). Averaging these standardized shapes provides an interpretable profile of group differences, with simplified burst characteristics. (Figure 2b). We capture the similarity between individual bursts by constructing the K-nearest neighbor (KNN) graph, linking each burst shape to its *K* most similar burst shapes, measured with the Wasserstein distance (also called Earth Mover’s distance) (Figure 2c, see Methods). From this graph, recording-level predictions were obtained by aggregating the neighbors of individual bursts (Figure 2d). We then applied spectral embedding to the KNN graph, preserving its structure while projecting burst shapes into a low-dimensional space (Figure 2e). Averaging the two-dimensional representation of all bursts in the recording yielded two shape summary features (Figure 2e). Finally, we combined these shape features with the traditional features in an XGBoost classification (Figure 2f), comparing the performance between classifiers. This representation of burst shapes provides the basis for evaluating their contribution to classifying experimental conditions, as detailed in the following section.

**Fig. 2.**
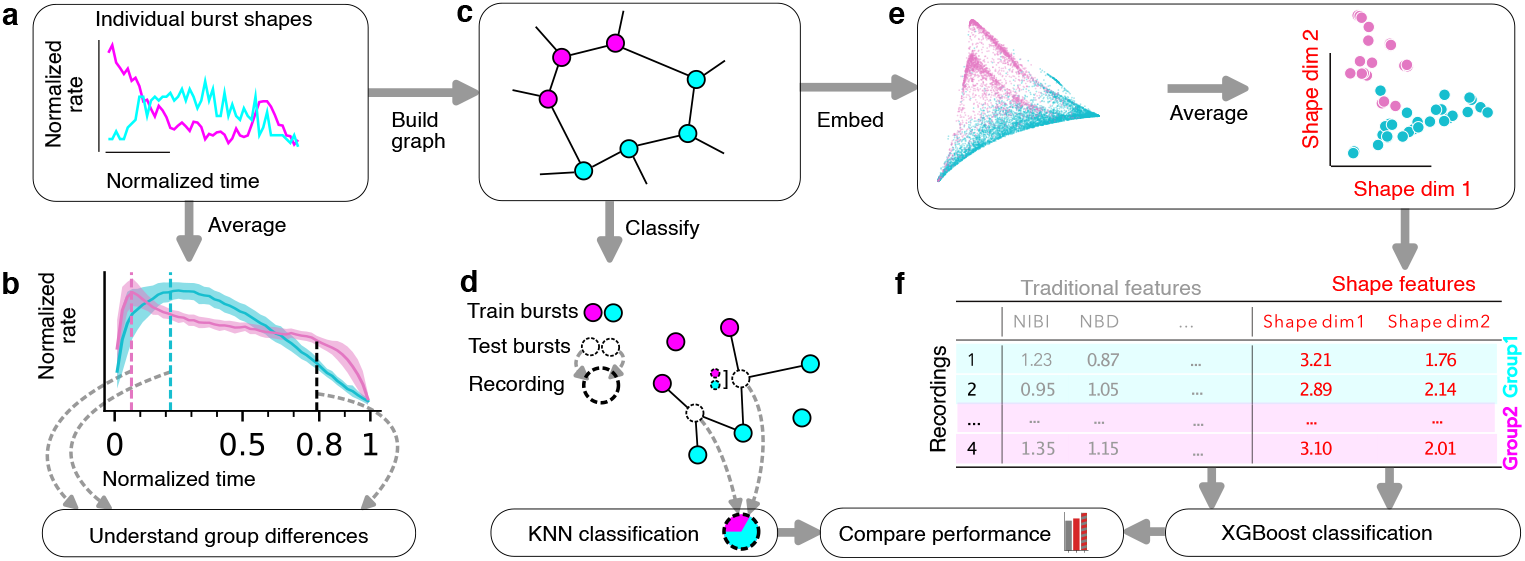
Framework for data-driven quantification of burst shapes. **(a)** Standardized burst shapes are computed by normalizing duration and size. **(b)** Understanding the group differences by averaging across recordings. **(c)** KNN graph connects each node (burst shape) to the *K* most similar nodes. **(d)** Classifying recordings based on individual burst shapes’ neighbors aggregated to the recording level. **(e)** Spectral embedding of individual burst shapes is aggregated (averaged) to the recording level, where the shape dimensions define the shape features. **(f)** Traditional and shape summary features are combined to predict recording identities using XGBoost.

### Augmenting traditional features with burst shape features improves classification performance across datasets

We compared the performance of four different classification models: KNN clustering using individual burst shapes (Figure 2d) and XGBoost classification using three feature sets: traditional, burst shape, and combined (Figure 2f), finding that including burst shape information significantly improved classifier performance (Figure 3). For evaluation, we employed an extensive cross-validation scheme, consisting of 100 repeated cross-validations with an 80-20 split, carefully correcting standard error of the mean for data interdependence [27, 28]. Generally, all models performed well above chance-level (Figure 3a).

**Fig. 3.**
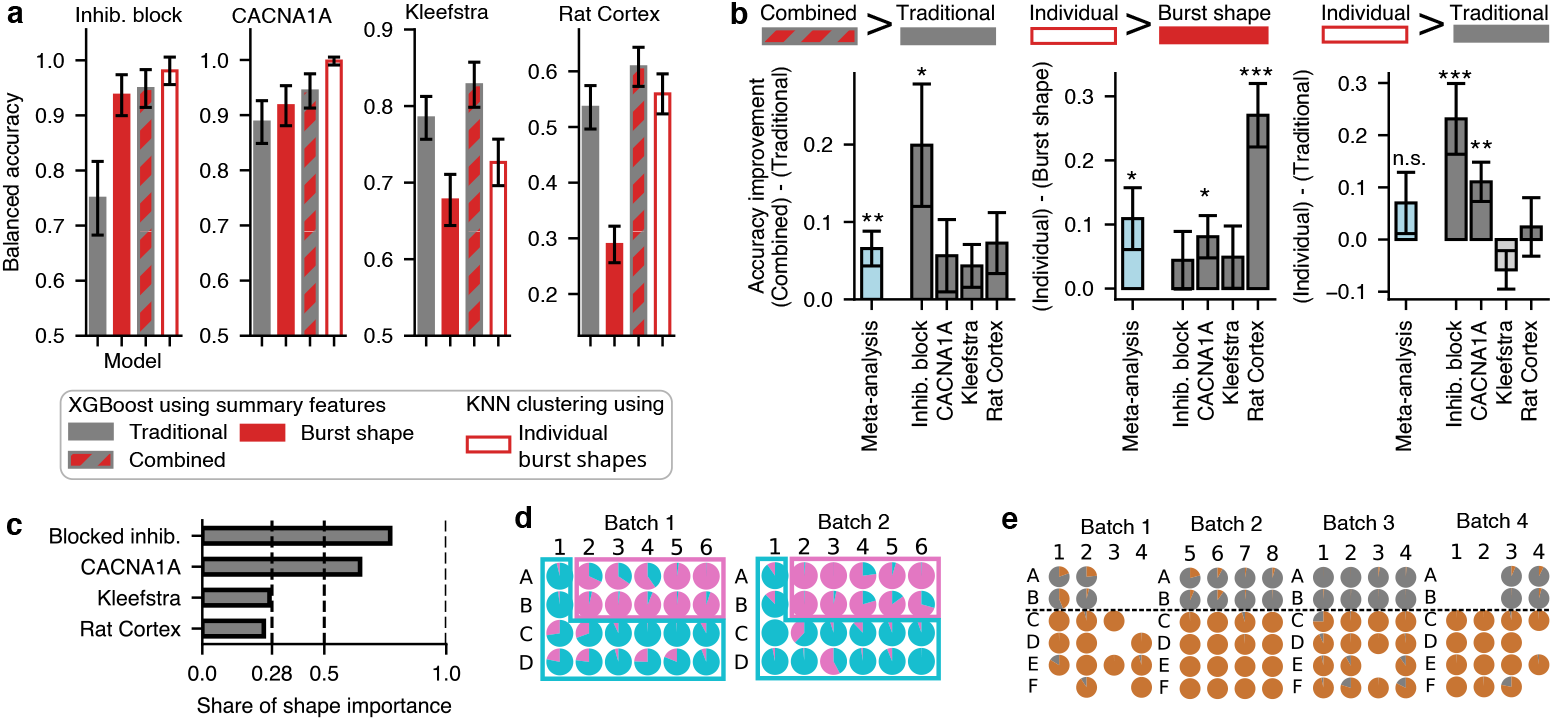
Augmenting traditional features with burst shape features improves classification performance across datasets. **(a)** Mean classification accuracies for four models: KNN clustering based on individual burst shapes, and XBoost models based on three summary feature sets (traditional, shape, and combined). Error bars, standard error of the mean; y-axis minima represent chance levels. **(b)** Difference in accuracies between models trained on datasets indicated in axis label. Combined features set outperforms the traditional set, individual burst features outperform shape summary features, and in some cases outperform traditional features. Error bars – standard error of the mean (n=100 shuffle splits, 80–20 ratio). Stars indicate significance; meta-analysis (hypothesis stated on top of each panel) used a random-effects model. **(c)** Shapley value analysis of the seven combined features shows that the two shape features substantially contribute to prediction. **(d)** KNN classification predictions (leave-one-out) for the blocked inhibition dataset in the layout with 4x6 wells per batch. Pie-chart shares represent the weighted neighbors of the recording; solid borders indicate the true class. **(e)** KNN classification predictions for the *CACNA1A* dataset. The dashed line separates the true labels (top: control, bottom: *CACNA1A*).

Pooling results across datasets in a meta-analysis showed that the combined feature set was significantly more accurate than the traditional features (Figure 3b, *p* = 3.4 *×* 10^*−*3^). On every dataset, using the combined features leads to an improvement that is larger than the standard error of the mean. In blocked inhibition, the balanced accuracy increased from 75% to 95%, improving from mediocre classification performance to an excellent classification (*p* = 1.3 *×* 10^*−*2^). To evaluate what drives the accuracy improvement of the combined feature set, we computed the feature importance of the seven combined features with Shapley values [29]. We exploited the additive nature of Shapley values, which allowed us to compute the relative share of the two shape features compared to the five traditional features (Figure 3c). This feature importance shows that in blocked inhibition and *CACNA1A* data, the burst shape features dominated the prediction with a share larger than 50%. For the Kleefstra and rat cortex data, the share is approximately similar to the share of features (2/7=28%). Burst shape features also rank high in the rankings of feature importance (Supplementary Figure S9b-g). This indicates that adding shape summary features to the traditional features consistently improves performance, with a substantial contribution of shape features.

Classification using individual burst shapes via the KNN graph is significantly more accurate than using burst shape summary features (Figure 3b, *p* = 2.3 *×* 10^*−*2^). On every dataset, the improvements are larger than the standard error of the mean, where improvements in the rat cortex (*p* = 3.5 *×* 10^*−*7^) and *CACNA1A* (*p* = 1.6 *×* 10^*−*2^) data are significant. This indicates that some information is lost when compressing the KNN graph with spectral embedding into shape summary features.

KNN-based classification using individual burst shape (Figure 2d) features is more accurate than using the traditional burst features for the blocked inhibition (*p* = 9.4 *×* 10^*−*4^) and the *CACNA1A* data (*p* = 4.1 *×* 10^*−*3^, Figure 3b). The generalized conclusion with meta-analysis is not significant because the traditional features perform better in the Kleefstra data, and the effect is unclear in the rat cortex data. This indicates that in some datasets, burst shapes alone can provide excellent predictions, while in other datasets, burst shapes can help by augmenting traditional features.

Using individual burst shapes in KNN classification achieves near-perfect accuracy in blocked inhibition (98%) and *CACNA1A* data (100%), reflected in the probabilistic predictions for each recording (Figure 3d, e). Most of the pie charts not only have a larger share for the correct class (meaning, correct prediction), but also are mostly filled, indicating that the algorithm also has a high confidence in those predictions.

Together, these results demonstrate that burst shapes contain substantial information about network phenotypes, especially in blocked-inhibition and *CACNA1A*. Thus, burst shapes are instrumental for describing network activity, complementing traditional features of bursting activity.

### Averaged normalized burst shapes reveal altered onset and offset dynamics

While spectral embedding produces compact features that perform well in classification, their abstract dimensions are not readily interpretable, providing only limited guidance for future mechanistic modeling. To address this, we sought simplified features from averaged burst shapes that capture biologically meaningful aspects of onset and termination dynamics while retaining predictive performance close to that of spectral embedding.

The normalized burst shapes can be directly averaged (Figure 2b), while raw burst shapes are difficult to compare due to varying sizes and durations. Indeed, the averages of normalized burst shapes (Figure 4a) have tight group means (±3 *×* standard error of the mean) compared with raw burst shape averages (Figure 4b). Large, long bursts dominate the raw burst averages, obscuring shape features. While differing onset dynamics can be detected from raw burst averages, the misaligned termination obscures the termination dynamics (Supplementary Figure S3). In contrast, the averages of normalized bursts (Figure 4a) preserve the timing of initiation and termination, and balance burst sizes, revealing consistent shape alterations.

**Fig. 4.**
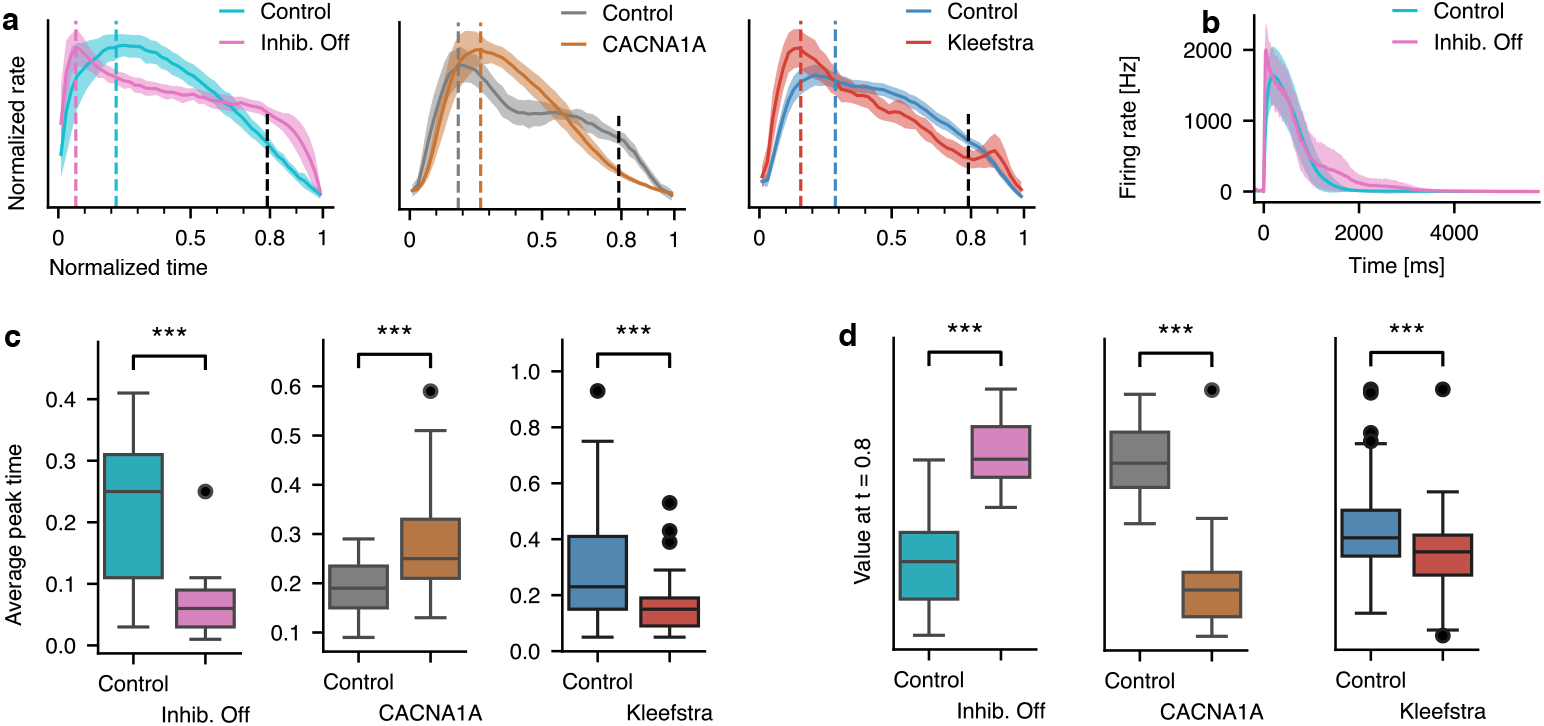
Averaged normalized burst shapes reveal altered onset and offset dynamics. **(a)** Averages of normalized bursts from control cultures and cultures with blocked inhibition, *CACNA1A* haploinsufficiency, and Kleefstra syndrome. Shades indicate ±3*×* standard error of the mean. **(b)** Averages of raw (unnormalized) burst shapes per recording are dominated by large bursts, obscuring the features evident after normalization (example of cultures with blocked inhibition). **(c)** Average peak timing (colored dashed lines in a) differs significantly from controls in all experiments, with earlier peaks in blocked-inhibition and Kleefstra, and later peaks in *CACNA1A*. **(d)** Amplitude near burst termination (80% of duration, black dashed line in a) is significantly higher for blocked inhibition, indicating abrupt termination, and reduced in *CACNA1A* and Kleefstra, consistent with prolonged decay dynamics. Stars indicate significance (*: *p <* 0.05, **: *p <* 0.01, ***: *p <* 0.001).

We characterized the initiation phase of bursts by the peak of activity (Figure 4c), similar to traditional approaches on raw burst shapes. Bursts in cultures with blocked inhibition reached the peak significantly faster compared to controls (inhib. off=0.06, control=0.21, *p* = 3.1 *×* 10^*−*7^), consistent with accelerated recruitment of network activity reported previously[24]. *CACNA1A* burst shapes showed a significantly delayed peak compared to control recordings (control=0.17, *CACNA1A*=0.26, *p* = 6.4 *×* 10^*−*6^), whereas Kleefstra syndrome average burst shapes peaked earlier than controls (control=0.28, Kleefstra=0.15, *p* = 7.7 *×* 10^*−*14^). Both findings are consistent with the original studies on raw burst shapes[9, 25] (also Supplementary Figure S3).

Termination dynamics were quantified by the activity level at 80% of normalized burst duration (Figure 4d), assessing whether bursts exhibit sustained activity with a sudden end or a prolonged decay. Cultures with blocked inhibition have a significantly higher activity near termination (t=0.8, Inhib. off=0.018, control=0.012, *p* = 4.0*×* 10^*−*12^), indicating that the lack of inhibition results in a sudden drop-off in activity. *CACNA1A* cultures display a significantly decreased level of activity before burst termination (t=0.8, *CACNA1A*=0.008, Control=0.016, *p* = 1.1 *×* 10^*−*17^), and Kleefstra syndrome displays a similar but smaller and significant reduction (t=0.8, Kleefstra=0.012, Control=0.015, *p* = 3.2 *×* 10^*−*5^), indicating more continuous, prolonged decay dynamics.

Our carefully designed onset- and offset features contain approximately the same amount of information as two spectral dimensions, but significantly less than the KNN-based clustering (Supplementary Figure S10). Thus, these burst shape features capture biologically meaningful aspects of the culture phenotype, useful for understanding the network dynamics and mechanistic modeling.

### Genetically related cultures exhibit larger burst shape similarity

Next, we asked whether genetic factors play an important role in shaping burst dynamics. To address this, we analyzed recordings from cultures with different levels of genetic relatedness and measured burst shape similarity using pairwise Wasserstein distances, comparing them to baseline distances from random pairs (Figure 5).

**Fig. 5.**
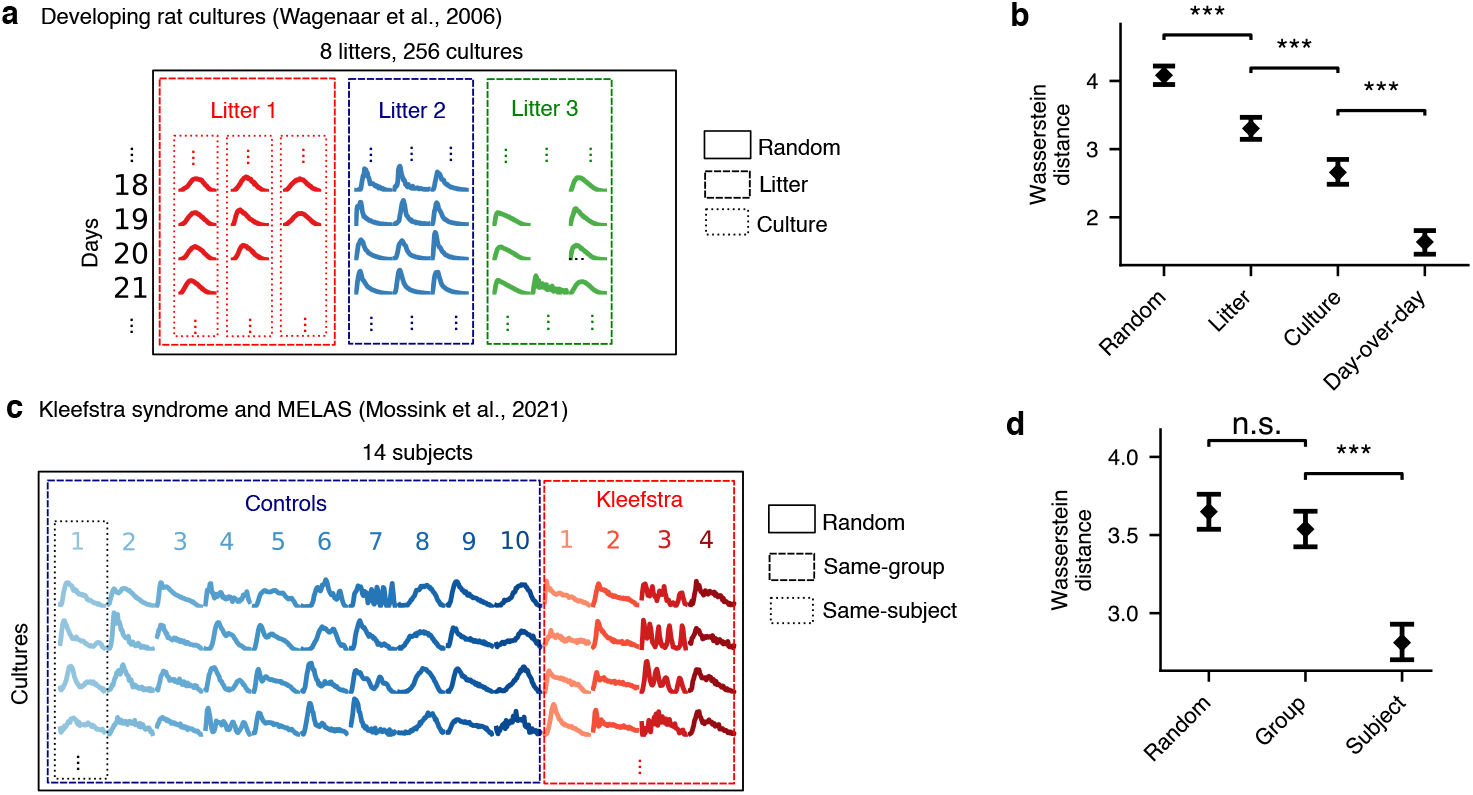
Genetically related cultures exhibit larger burst shape similarity. **(a)** Example average burst shapes per recording of rat primary cultures during development [15] are grouped across litters and cultures. Frames indicate comparison groups for panel (b). **(b)** Average Wasserstein distances between bursts show progressively increasing similarity: pairs from the same litter are closer than those from different litters, pairs from the same culture are closer than those from different cultures in the litter, and day-over-day variation within a culture is the smallest of all. **(c)** Average burst shape examples from the hPSC disease dataset [9]. **(d)** Recordings from the same subject have significantly smaller burst shape distances, while the disease group has no significant effect. Error bars represent the confidence interval. Stars indicate significance (*: *p <* 0.05, **: *p <* 0.01, ***: *p <* 0.001).

In developing rat primary cultures [15], we found that burst shapes were stable across litters and throughout development. Recordings from the same litter (batches) exhibited significantly smaller burst shape distances than random comparisons (two-sided t-test, *p* = 3.0 *×* 10^*−*12^, Figure 5c). Distances within individual cultures measured across all recording days were even smaller (*p* = 3.1 *×* 10^*−*7^), and day-to-day variability within a culture was the smallest of all (*p* = 7.1 *×* 10^*−*14^). These findings indicate that each culture develops a characteristic burst shape that changes only slowly over time. The strong within-litter similarity suggests a genetic component, although other factors, such as seeding conditions, may also contribute.

In hPSC-derived cultures [9], we found that burst shapes were strongly influenced by subject-specific genetics. Recordings from the same subject displayed significantly smaller distances than random comparisons (*p* = 4.3 *×* 10^*−*17^, Figure 5d). The disease status did not significantly influence the average burst similarity; however, as we have seen above (Figure 3a, 4c,d), the identity of the most similar bursts still carried information about the group identity. These results together indicate that both disease status and genetics influence the similarity of the burst shapes.

Overall, our findings demonstrate that burst shapes are stable across development and reflect genetic identity in both rat primary and hPSC-derived neural cultures.

## Discussion

Network bursts in neuronal cultures and during development *in vivo* exhibit a wide variety of profiles and durations, making their burst shapes difficult to quantify systematically. In this work, we present a data-driven approach to solving this problem, yielding a framework for extracting meaningful and practical descriptions of burst shapes. We normalize bursts to a common duration and size, enabling the pooling of variable burst shapes and the use of distance-based metrics to capture information in their profiles. This approach enhances the phenotyping of networks by using burst shape information without the need to extract specific descriptors. Thus, burst shape features contain sufficient information to distinguish between experimental conditions, such as pathological gene variants or pharmacological perturbations of neuronal activity. Moreover, we show that burst shapes also reflect individual genetic variance carried by cultured neurons.

Our approach is data-driven, does not require model selection, and still yields interpretable features that allow for direct examination of burst dynamics. Normalized burst shapes can be directly visualized and compared. For instance, when we analyzed burst shapes from cultures with blocked inhibition, we observed that both the onset and offset of bursts became faster. The faster onset dynamics are consistent with previous work on inhibition blockade [24], but we additionally emphasize offset dynamics, which are typically obscured when bursts are averaged directly. Interestingly, such changes in the offset had been predicted by mechanistic models of network bursting activity [18]. Similarly, we observed that *CACNA1A* haploinsufficiency prolonged burst onset and strongly smoothed burst offset. Both effects align with changes associated with *CACNA1A* loss-of-function and compensatory mechanisms [25]. While the first effect confirms what was observed in the original publication, the offset change is novel, and its effect size appears to be even stronger. Overall, this implies that burst shapes can serve as a novel and sensitive indicator of proximity to a healthy state. As such, it can be further used in pharmacological or gene therapy experiments aimed at understanding how new or existing drugs can correct the effects observed in patient cell lines [30–33].

Variance in bursting activity across cultures and experiments is often considered to be large. There is extensive evidence that each culture may have a unique activity profile. Upon stimulation, cultures can show individually specific patterns of activity [34]. At the same time, early experimental work by Wagenaar et al. [15] showed that part of this variance stems from genetic differences. Specifically, they demonstrated that “sister cultures,” cultures derived from the same litter, have lower variability compared to non-sister cultures [15]. In our work, we reanalyzed their data and found that burst shapes are also more similar when cultures originate from the same litter. Extending these findings, when we analyzed bursts from hPSC-derived cultures obtained from the same cell line (thus sharing the entire genome), we found that cultures with identical genomes exhibit more similar burst shapes. This emphasizes that burst shape may be a particularly useful readout for studying genotype–phenotype associations in *in vitro* systems, although the precise mechanisms underlying this association remain to be determined.

Our approach relies on accurate burst detection, as errors in defining burst start and end points can misalign shapes and obscure averages. In this study, a simple detection algorithm with interactive review was sufficient, but more complex datasets may require more sophisticated detection or alignment. Normalization and embedding choices also influence the framework. While spectral embedding proved effective here, alternative methods (e.g., higher-dimensional spectral spaces or graph-embedding algorithms such as node2vec) could capture additional structure. Finally, we collapsed spiking activity across electrodes into a single time series, which facilitates application to high-throughput MEA systems but ignores spatial propagation of activity. Incorporating two-dimensional burst dynamics, especially with high-resolution recordings [35–37], may provide further insights into variability and network structure.

Overall, we show that normalizing and comparing burst shapes provides additional information about spontaneous activity in developing neuronal cultures and improves the classification of dynamics. This opens new avenues for more detailed characterization of network phenotypes, particularly in the context of hPSC modeling of neuropsychiatric disorders, where changes in patient-derived cultures may be subtle.

## Methods

### Data

We utilized four distinct micro-electrode array (MEA) datasets from neural cultures, including both rat and human pluripotent stem cell (hPSC)-derived cultures. Two of these datasets were publicly available, while the other two were provided by collaborators. For all datasets, we used the processed spiking data, which consists of the timestamped spike trains for each electrode.

### Blocked inhibition

This dataset was provided by Vinogradov et al. [26] and consists of recordings from dissociated E18 mouse cortical neurons. The neurons were cultured on 24-well MEA plates, each well containing 12 gold electrodes. For two batches, the spontaneous network activity was recorded at 17 and 18 days in vitro (batches 1 and 2). To block synaptic inhibition, 40*µ*M of Bicuculline was added to 10 of the 24 wells in each batch. Following a 10-minute stabilization period, network activity was recorded for 20 minutes. While the original study did not discuss burst shapes, this dataset is well-suited for our analysis due to its systematic modulation of network activity.

### *CACNA1A* haploinsufficiency

This dataset was provided by Hommersom et al. [25] and consists of human induced pluripotent stem cell (iPSC)-derived cultures. The control line was modified with CRISPR/Cas9 to generate *CACNA1A* haploinsufficiency. The cultures were recorded for 5 minutes using 48-well MEA plates, each well containing 16 electrodes. The data comprises four batches, with 24 recordings from *CACNA1A* cultures and 57 controls (not all wells were used for that study). The original study reported a slightly delayed peak time of bursts from *CACNA1A* haploinsufficiency compared to controls, which suggests a systematic burst shape difference relevant to our investigation.

### Kleefstra syndrome

This dataset was published by Frega et al. [13] and consists of induced hPSC cultures derived from skin fibroblasts of control and disease patients. The dataset includes cultures from four Kleefstra patients, five healthy controls, and five isogenic controls (modified patient cells). The study also contained three MELAS patients, which we excluded from our analysis, because the original study reported that MELAS burst shapes were very similar to those of controls. In contrast, Kleefstra bursts were reported to have differing onset dynamics, making them suitable for our investigation and simplifying the classification to a binary problem of Kleefstra versus control networks.

### Developing embryo rat cultures

This dataset was published by Wagenaar et al. [15] and consists of anterior cortical neural cultures from Wistar rat embryos, extracted on day 18 of gestation. Cells from one litter were combined, and in total, eight batches (litters) were collected. Eight batches were collected, where each batch is from a single litter, i.e., a mixture of several embryos with shared genetics. The cultures were recorded for 30 min without a prior stabilization period. We selected the data between days 7 and 35 in vitro for our analysis. While focusing on developmental trajectories of burst summary statistics, the original study also included simple features of intra-burst dynamics like the onset and offset durations. They reported that burst durations decreased over development, in particular, the burst onset duration shortened, see Figure 6 in the original study.

### Preprocessing

#### Burst identification

To identify network bursts, we first concatenated the spike times from all recording units (electrodes) into a single network-wide spike train. We then used a detection algorithm to find the start and end of bursts based on the inter-spike interval (ISI). A burst was defined to start when the inter-spike interval fell below a predefined threshold (*maxISIstart*) and to end when the next ISI exceeded a second threshold (*maxISIb*). While these thresholds were often identical, they could be set independently of each other.

To filter for meaningful network events, we applied additional criteria. A detected burst was accepted as a burst only if it (1) had a minimum burst duration(*minBdur*), (2) a minimum inter-burst interval (*minIBI*), and (3) a minimum number of spikes (*minSburst*). We also applied optional filters for a minimum burst length and a minimum firing rate. In some instances, a 4 ms smoothing kernel was also applied to mitigate the influence of fluctuations in very short bursts.

The chosen parameters are detailed in Supplementary Table S2.

#### Burst review

After running the automated detection algorithm, we manually reviewed the burst detection using a custom interactive tool (Supplementary Figure S1). This tool allowed us to efficiently examine the quality of burst detection across a large number of recordings. The tool overlays the detected bursts on traces of the firing rate and raster plots, with the option to zoom in for closer inspection. If the initial parameters did not accurately capture the network bursts, we could adjust the burst detection parameters until an acceptable level of accuracy was achieved.

### Spectral embedding of the burst shapes with Wasserstein distance

#### Normalization

To compare bursts using the Wasserstein distance, we normalized the bursts in both time and amplitude, thereby isolating their shape feature. We normalized duration to a length of 1, discretizing it into 50 time bins and calculating the firing rate for each bin from the spike times. Finally, we normalized each burst shape by its integral (the sum of all bins) to ensure a total integral of 1. These steps resulted in a standardized burst shape with unit duration and unit integral, which was then used in further analysis.

#### Wasserstein distance

To quantify the similarity between burst shapes, we chose the Wasserstein distance metric (also called Earth Mover’s distance). This metric was selected because it is robust to small shifts, assigning a small distance to similar burst shapes and a large distance to dissimilar ones. For two bursts *b*_1_ and *b*_2_, the one-dimensional Wasserstein distance *d*(*·, ·*) can be calculated from their cumulative sums:

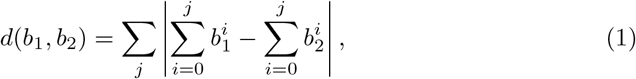

where 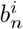 is the value of burst *n* at bin *i*.

#### K-nearest neighbors (KNN) graph

To capture the relationships between individual burst shapes, we constructed a KNN graph, which connects each burst shape with its *K* most similar burst shapes. The pairwise distances between all burst shapes were computed using the Wasserstein distance. The number of nearest neighbors *K* was selected such that each node was connected to approximately 1% of all nodes (similar to the recommendation for t-SNE perplexity by Kobak and Berens [38]), resulting in *K* between 85 and 150 for our datasets. This KNN graph served as the basis for both KNN clustering and spectral embedding.

#### Spectral embedding

To obtain a low-dimensional embedding from the KNN graph, we performed spectral embedding. First, we constructed the symmetric adjacency matrix 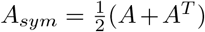 from the unsymmetric adjacency matrix *A* of the KNN graph. Next, we computed the Graph Laplacian *L* = *D – A*_*sym*_, where *D* is the degree matrix such that each row (and column) of *L* sums up to zero. Then, we used the eigenvectors of *L* corresponding to the smallest non-zero eigenvalues as the low-dimensional coordinates, as they best preserve the connectivity structure of the underlying graph.

### Classification of recordings

To understand which burst features contain biologically meaningful information, we classified recordings with various models and feature sets. We employed two primary classification strategies. First, we used KNN-based classification to leverage the rich information within individual burst shapes, aggregating this information to the recording level. Second, we applied XGBoost to classify recordings based on summary features, including traditional, shape, and combined feature sets. For all classification tasks, we evaluated model performance using balanced accuracy and a repeated stratified shuffle split (100 times, 80-20 training-testing split) for robust cross-validation.

#### KNN-based classification leveraging individual burst shapes

To leverage the rich information present in individual burst shapes, we performed a KNN-based classification. To prevent data leakage, we first ensured that no individual bursts from test-set recordings were included in the training set. Then, for each of these ‘test bursts’, we identified its *K* nearest neighbors among the ‘training bursts’. We predicted the class of the ‘test burst’ with a voting scheme based on these neighbors, which belonged to a training recording with a known class label. To account for class imbalances and recording-specific biases, we weighted the vote of each neighbor. The weight was calculated as the inverse of the number of bursts in its recording 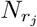 and the inverse of the number of recordings in its class 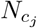.

The class vote vector **v**_*i*_ for an individual burst was computed as

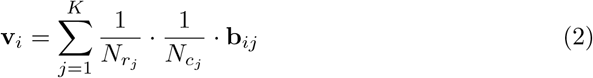

where **b**_*ij*_ is a one-hot vector indicating the class label of the *j*-th neighbor of test burst *i*. To obtain a prediction for a test recording, we averaged the vote vectors **v**_*i*_ across all its constituent bursts. The class with the highest average vote was assigned as the predicted label for that recording.

#### XGBoost prediction from recording summary statistics

To assess the predictive power of shape features relative to traditional burst features, we trained XGBoost on three sets of recording-level summary statistics: shape features, traditional features, and a combined feature set.

The *traditional burst features*, used as a baseline for prediction accuracy, were derived from commonly used burst metrics[10]:

- Mean firing rate (MFR): average of the culture’s firing rate
- Burst spike rate (BSR): average of the bursts’ firing rates
- Network burst rate (NBR): number of bursts divided by recording duration
- Network burst duration (NBD): average of burst durations
- Percentage of Random Spikes (PRS): percentage of spikes not assigned to bursts

We excluded some features used by van Hugte et al. [10] (mean burst rate (MBR), burst duration (BD), and high-frequency bursts (HFB)) because they rely on single-channel bursts and high-frequency bursts, which were not computed in our analysis.

Our *burst shape features* were based on the spectral embedding, the low-dimensional projection of the KNN graph. For each recording, we averaged the individual burst shape locations in two dimensions. This yielded two shape summary features per recording (spectral embedding dimensions 1 and 2). While more or fewer dimensions than two were possible, we always chose two dimensions, the same as in the visualizations. The *combined feature set* is obtained by combining the five traditional and the two burst shape features.

For all XGBoost models, we employed a nested cross-validation approach with an outer (testing) and inner loop (validation). The outer loop consisted of a repeated stratified shuffle split (100 times, 80-20 split) for testing (same as in KNN classification). The inner loop optimized the XGBoost hyperparameters via a grid search in a 5-fold cross-validation. We report the balanced accuracy on the test set to evaluate the predictive power of each feature set.

#### Feature importance with Shapley values

To quantify the relative contribution of traditional and shape features, we computed Shapley values [29] (feature importance) for the XGBoost model trained on the combined feature set. Shapley values are a robust method from cooperative game theory for fairly distributing the ‘payout’ (in this case, the model’s prediction) among the ‘players’ (the features). This approach is used in Machine Learning to quantify the importance of features while accounting for complex interactions. A particularly useful property of Shapley values is their additivity, which allowed us to calculate the total importance of the traditional feature set and directly compare it to that of the shape features.

### Statistics

#### Comparison of model accuracies

For comparing model accuracies, we recognized that standard methods such as a t-test lead to inflated significance because accuracies from cross-validation are interdependent due to the reuse of data [27, 28]. Therefore, to robustly compare model accuracies, we used the correction for the standard error of the mean 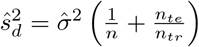, where *n* is the number of trials, *n*_*te*_ is test set size, and *n*_*t*_*r* is training set size. With our cross-validation parameters (*n* = 100 and 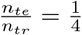), we computed a paired two-sided t-test. To test the overall statement across datasets, we computed a meta-analysis using a random-effects model.

#### Distance comparison

We analyzed the distances between recordings using a multi-step process. First, we calculated the pair-wise distances between recordings by averaging the distances between their constituent individual burst shapes. We then filtered the pairwise distances based on specific criteria, for example, including all distances (“ random”) or only distances between recordings from the same subject (“ subject”). To balance the dataset and avoid falsely inflating sample size, we computed a single average distance for each recording by averaging all outgoing distances. Finally, we used the two-sided t-test to determine if there were significant differences in these average distances.

## Data and code availability

- All code is publicly available on GitHub https://github.com/LevinaLab/burst-clustering.
- Wagenaar et al. [15], and Mossink et al. [9] data is publicly available from the respective publications.
- We make the Vinogradov et al. [26] data publicly available with this publication.
- Data from Hommersom et al. [25] is available from the original authors upon reasonable request per the original publication.

## Acknowledgements

This work was supported by a Sofja Kovalevskaja Award from the Alexander von Humboldt Foundation. TS received funding through the Else Kröner Medical Scientist Kolleg “ClinBrAIn: Artificial Intelligence for Clinical Brain Research”, and the Deutsche Forschungsgemeinschaft (DFG, German Research Foundation) under Germany’s Excellence Strategy – EXC number 2064/1 – Project number 390727645. TS and EG thank the International Max Planck Research School for Intelligent Systems (IMPRS-IS) for support, and OV thanks the International Max Planck Research School for the Mechanisms of Mental Function and Dysfunction and the Joachim Herz Foundation for support. We acknowledge the support from the BMBF through the Tübingen AI Center (FKZ: 01IS18039B). AL is a member of the Machine Learning Cluster of Excellence, EXC number 2064/1 – Project number 39072764.

We thank all lab members, particularly Aaron Spieler and Elias Seiffert, for helpful discussions and their support. We also thank Farzin Negahbani for helpful discussions.

A big thank you also to Marina Hommersom and Nina Doorn for sharing their data and helpful discussions.

## Supplemental information

**Table S1.** Summary statistics of datasets used in the analysis.

| Dataset | #cultures | #bursts | Time (min) | Total time (min) |
| --- | --- | --- | --- | --- |
| Blocked inhibition | 48 | 8571 | 20 | 960 |
| *CACNA1A* | 81 | 2114 | 5 | 405 |
| hPSC disease | 390 | 9350 | 20 | 7800 |
| Developing rat | 451 | 11914 | 60 | 27060 |

**Table S2.**
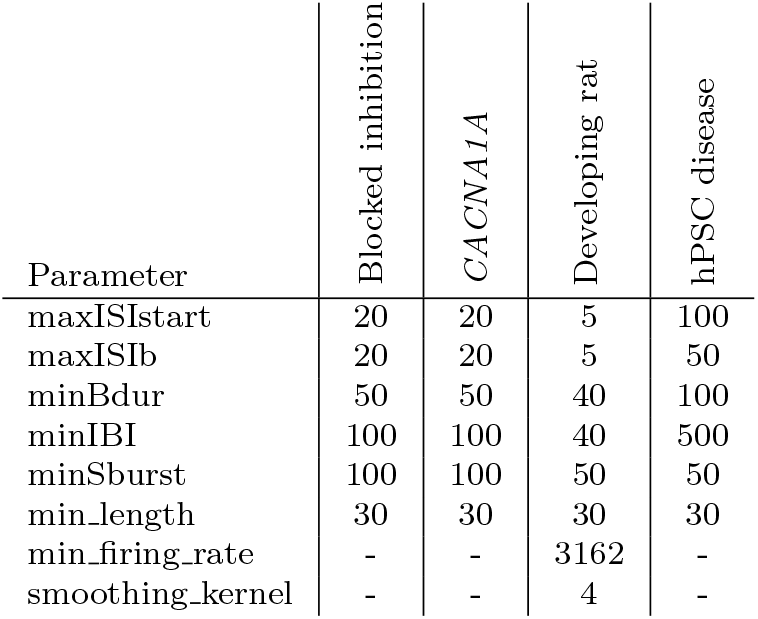
Network burst detection parameters. For blocked-inhibition (Bicuculline) [26], *CACNA1A* haploinsufficiency[25], developing embryo rat cultures [15], and hPSC disease recordings[9], we detect the bursts based on the parameters given in the table in milliseconds. ISI – inter-spike interval, maxISIstart – maximal ISI, maxISIb – minimal ISI during the burst, minBdur – minimal duration of the burst, minIBI – minimal inter-burst interval, minSburst – minimal number of spikes in burst, min length – minimal length of the burst, min firing rate – minimal average firing rate, smoothing kernel – length of smoothing kernel.

**Table S3.** Mean classification accuracies. Balanced accuracies for the cross-validated classification task across datasets. Traditional (baseline, highlighted in blue): XGBoost on summary features, shape: XGBoost on shape features, combined: both feature sets. KNN classification uses individual burst shapes aggregated to the recording level. Stars indicate frequentist comparison to the baseline (corrected for data interdependence):*: *p <* 0.05, **: *p <* 0.01, ***: *p <* 0.001. Daggers indicate Bayesian comparison to baseline: *†* moderate (posterior *P >* .7) evidence, *‡* strong (*P >* .9), (triple-dagger) very strong evidence (*P >* .99). Bold: best model.

| Dataset | Random | Summary Features |  |  | Individual burst shapes |
| --- | --- | --- | --- | --- | --- |
|  |  | Traditional | Shape | Combined |  |
| Blocked inhibition | 0.5 | 0.750 | 0.937 | 0.949* ‡ | **0.981\*\*\*** ‡ |
| *CACNA1A* | 0.5 | 0.887 | 0.917 | 0.944 † | **0.998\*\*** ‡ |
| hPSC disease (Kleefstra) | 0.5 | 0.785 | 0.677 | **0.828** † | 0.726 ‡ |
| Developing rat | 0.125 | 0.535 | 0.289 | **0.608** † | 0.559 |

**Fig. S1.**
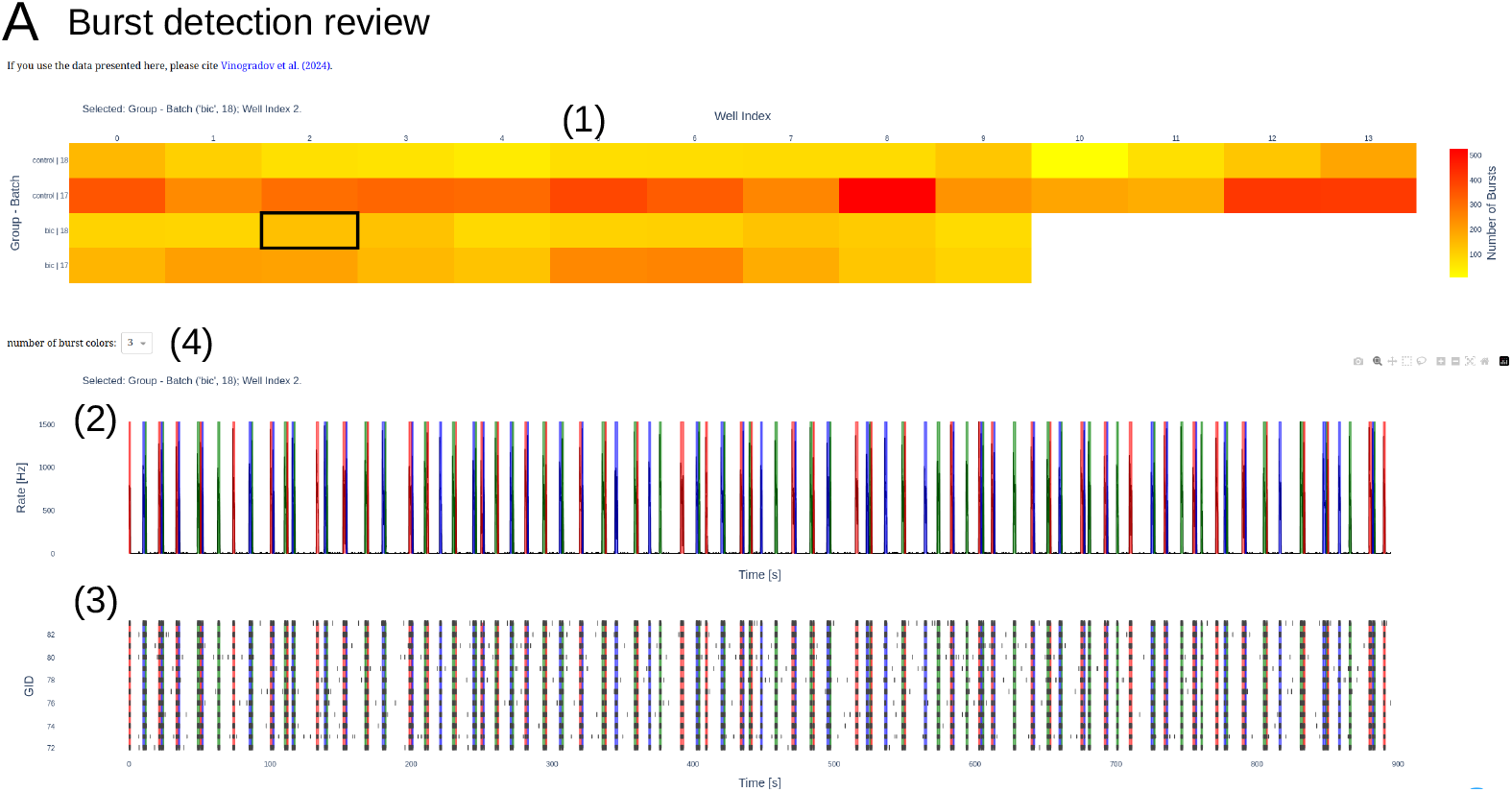
Interactive tools enable efficient processing of large amounts of data (part 1). Examples are shown for blocked inhibition data[26]. These tools can be deployed offline or online. **(A) Burst detection review:** This tool enables fast and detailed evaluation of the burst detection algorithm. We use it to improve the burst detection parameters iteratively. (1) Overview of all recordings. Rows and columns are the index: Here, it is Well Index for columns and Group-Batch for rows. The color indicates the number of bursts detected in the recording. Clicking on one of the recordings opens up the firing rate (2) and raster plot (3) for that recording. The firing rate and raster plot have a synchronized time axis. The detected bursts are highlighted with a colored background, where the number of colors can be adjusted for improved visualization (4). To inspect details of burst detection, zoom in by selecting a time frame. For some of the datasets presented here, the tools with the data are available online. Find the link through the GitHub repository or contact the first author.

**Fig. S2.**
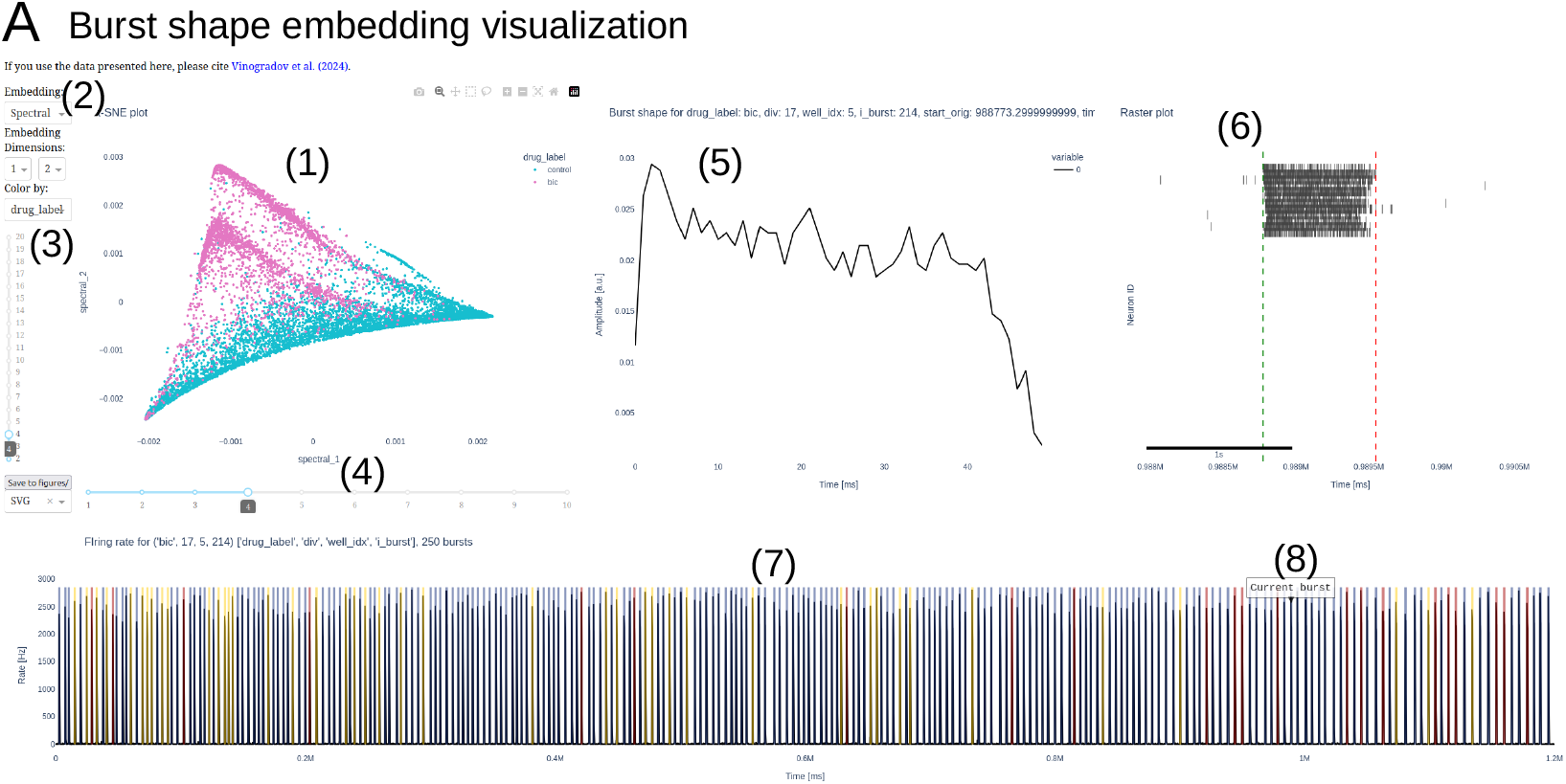
Interactive tools enable efficient processing of large amounts of data (part 2). Examples are shown for blocked inhibition data[26]. These tools can be deployed offline or online. **(A) Burst shape embedding visualization:** This tool enables fast interactive exploration of the embeddings of burst shapes. (1) The whole embedding of individual burst shapes can be explored by zooming in and selecting and deselecting certain groups. (2) Options to change the embedding, i.e., to PCA or t-SNE, to change the plotted dimensions (default is up to 20th dimension). There is also the possibility to colorcode the embedding by other criteria: batch, firing rate, burst duration, etc. One particular option, ‘cluster’, is the spectral clustering for which the number of clusters can be changed with the slider (4). When selecting (clicking) one of the nodes (burst shapes) in the embedding (1), the application opens the burst shape (5), the raster plot (6), and the associated recordings’ firing rate (7). The other bursts in the recording are highlighted with a background color, with the current burst especially highlighted (8). It is possible to directly switch to another burst from the same recording by clicking on it. The background color corresponds to the spectral clusters which can be changed with the slider (3). For some of the datasets presented here, the tools with the data are available online. Find the link through the GitHub repository or contact the first author.

**Fig. S3.**
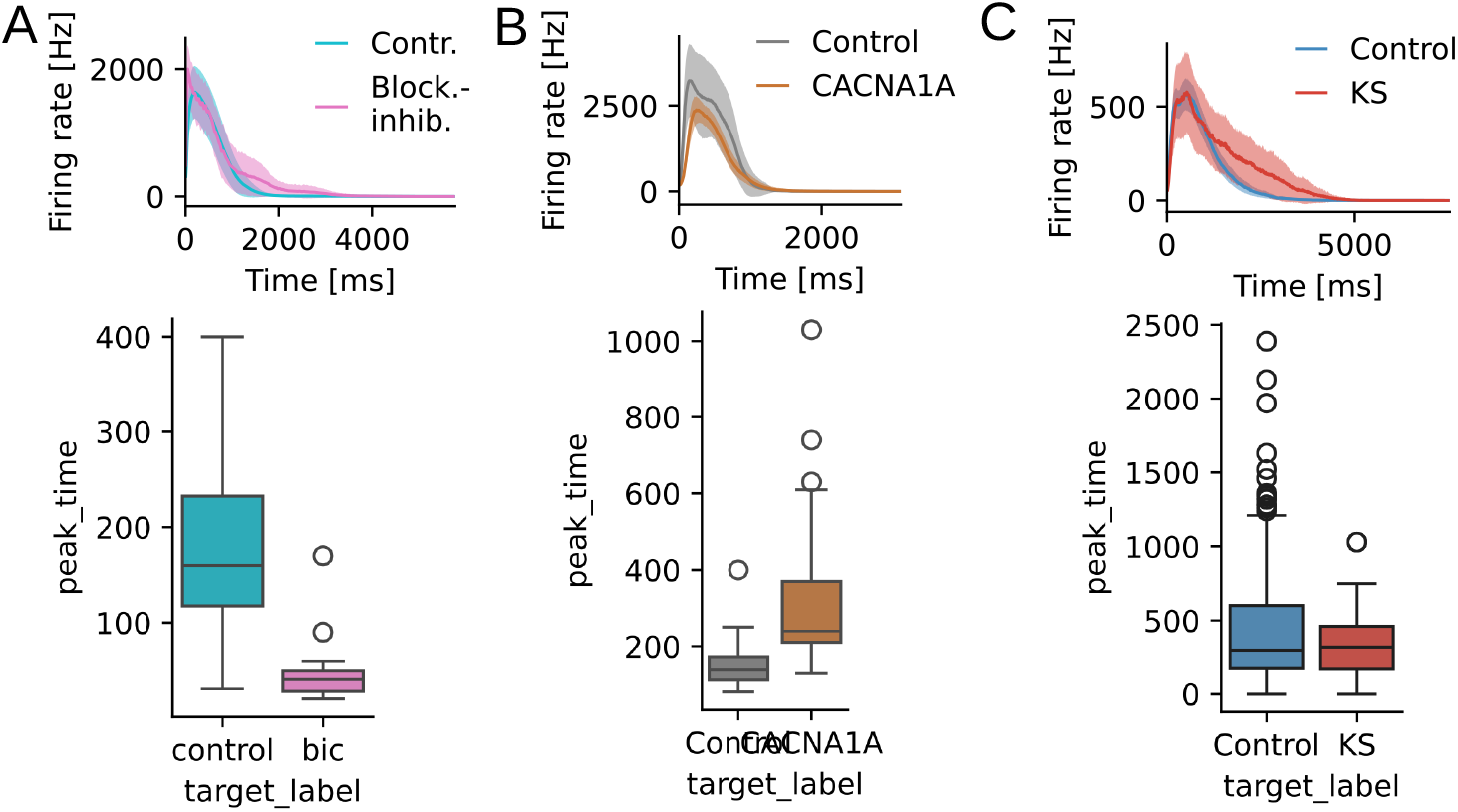
Raw shape differences between groups. Averaging raw burst shapes enables comparison of onset dynamics, but makes it impossible to quantify burst termination dynamics. **(A)** Blocked inhibition recordings burst shapes peak significantly earlier than controls (bic=46.5 ms, control=175.4 ms, p=1.1e-17). **(B)** *CACNA1A* recordings burst shapes peak significantly later than controls (control=155.0 ms, CACNA1A=298.6 ms, p=2.4e-7 **(C)** Kleefstra recording burst shapes peak significantly earlier than controls (control=445.5 ms, 344.1 ms, p=5.7e-3).

**Fig. S4.**
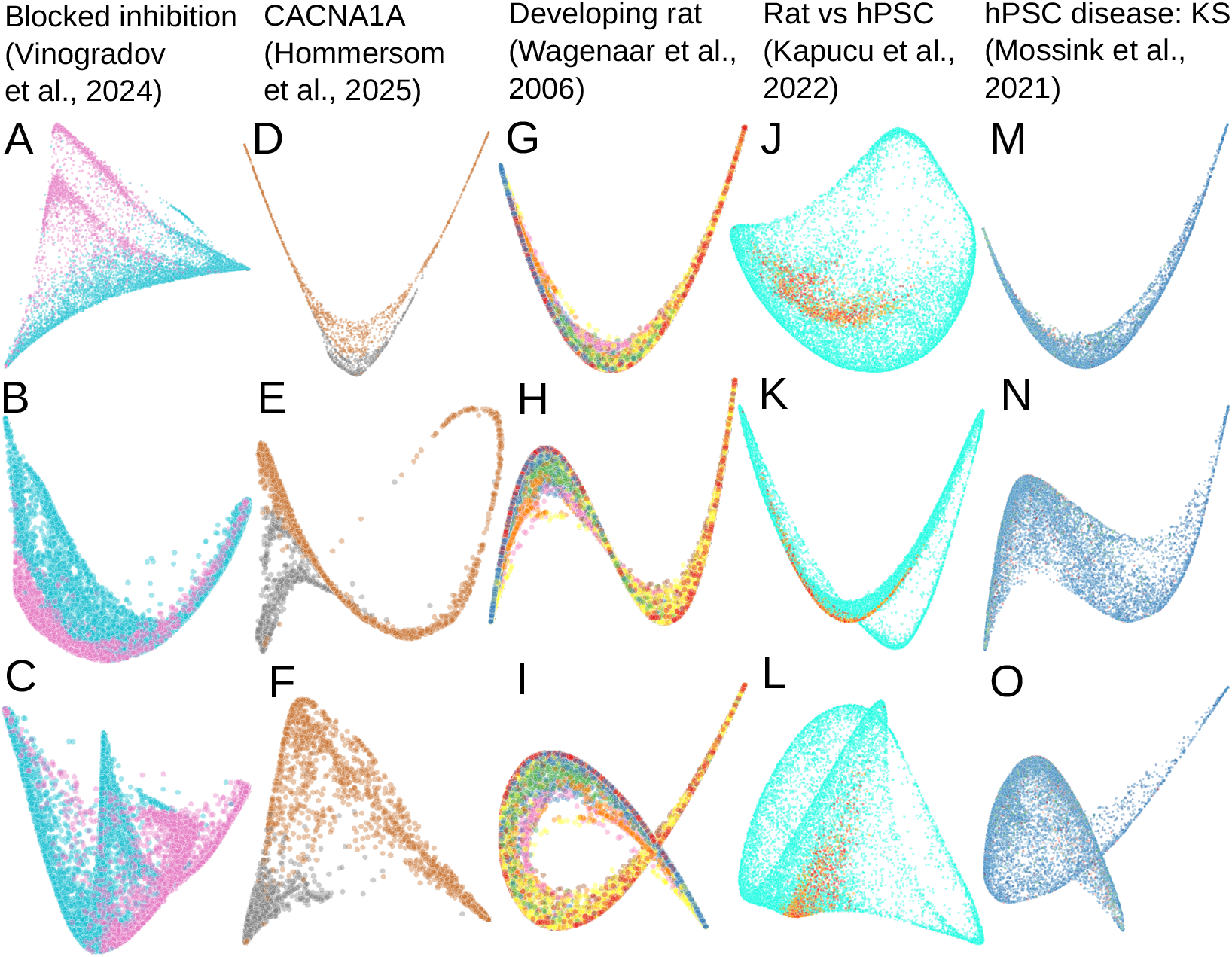
Spectral embeddings of the five datasets (dimensions 1 to 3). **Dimensions**: First row is first vs. second dimension. Second row is first vs. third dimension. Third row is second vs. third dimension. **Datasets:** **A-C** Blocked inhibition[26] **D-F** *CACNA1A* haplosufficiency insufficiency[25] **G-I** Developing rat primary cortex[15] **J-L** Rat vs. hPSC [39] **M-O** hPSC diseases: Kleefstra and MELAS[9] Note that some dimensions separate the groups very well, e.g., *CACNA1A* and control are well-separated in the second and third spectral dimension **(F)**.

**Fig. S5.**
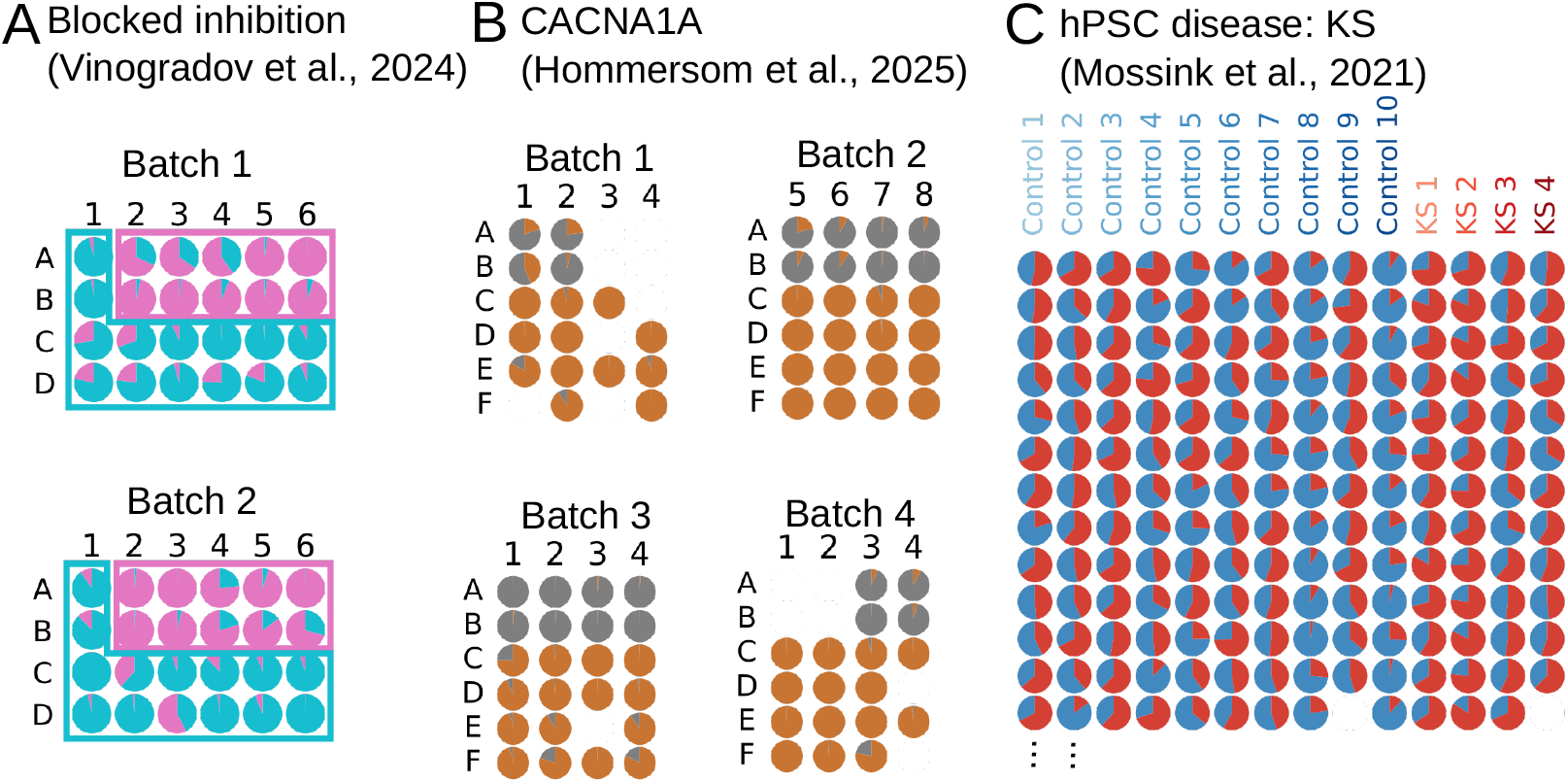
K-nearest neighbor classification predictions (part 1). Predictions are the *votes* from the K nearest neighbors weighted by class and number of bursts per recording. Each pie chart represents the share of these votes for one recording, indicating the confidence of the prediction. These predictions are obtained using leave-one-out cross-validation, in contrast to the main text, which employs a stricter cross-validation with a repeated 80-20 split. **(A)** Blocked inhibition[26]predictions plotted in the plate layout in which the cultures were recorded. Rectangles indicate the true labels. **(B)** *CACNA1A* haploinsufficiency vs control[25] predictions plotted in the plate layout in which the cultures were recorded. *CACNA1A* is located in rows A and B; controls are located in rows C-F. **(C)** Subset of hPSC disease recordings [9] (only Control vs Kleefstra) predictions sorted by subject.

**Fig. S6.**
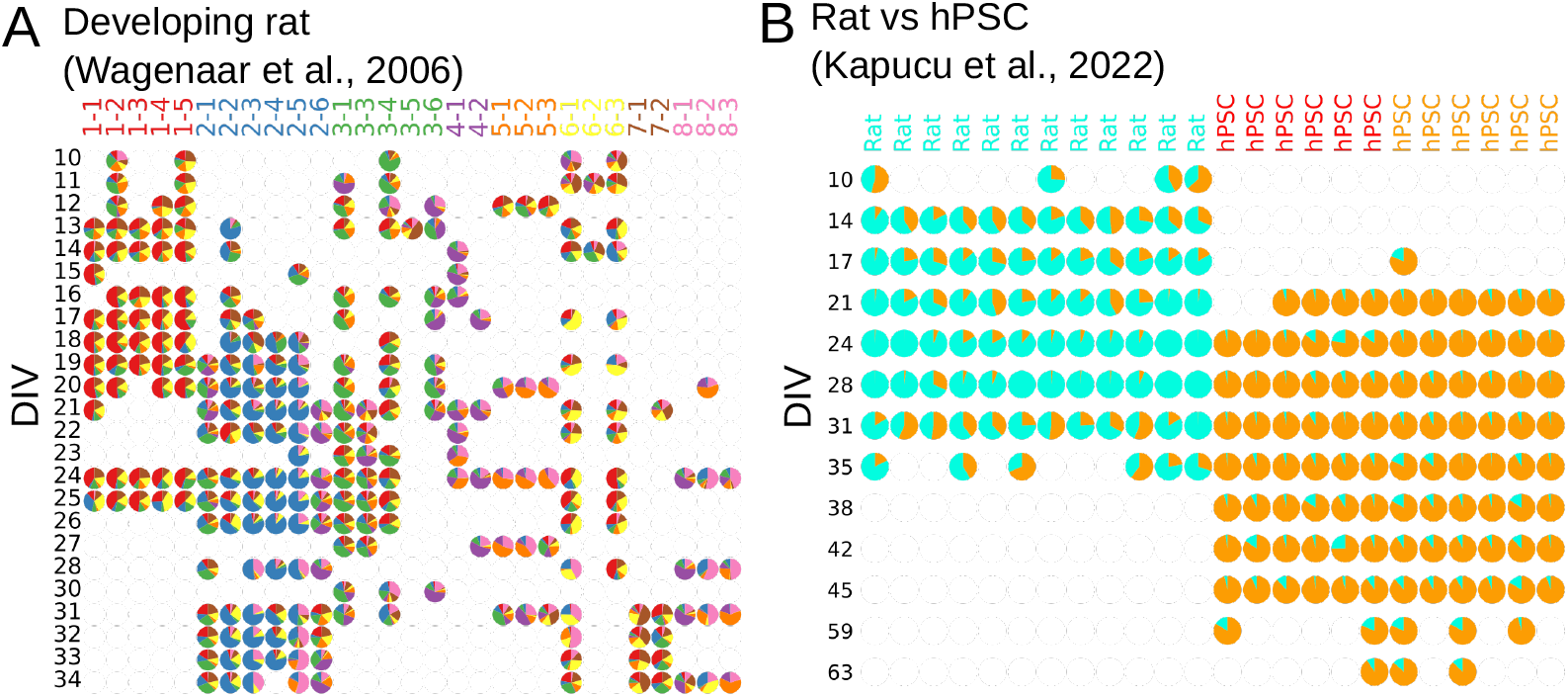
K-nearest neighbor classification predictions (part 2). Predictions are the *votes* from the K nearest neighbors weighted by class and number of bursts per recording. Each pie chart represents the share of these votes for one recording, indicating the confidence of the prediction. These predictions are obtained using leave-one-out cross-validation, in contrast to the main text, which employs a stricter cross-validation with a repeated 80-20 split. Columns are the days in vitro (DIV). Thus, developmental trajectories are from top to bottom. **(A)** Developing rat primary[15] recordings predictions. Batches (true labels) are the colors of the columns. Column labels are `[batch]` -`[index of culture within batch]`. **(B)** Developing Rat vs hPSC [39] recordings. Type of recording (true labels) is the colors of the columns. The two batches of hPSC recordings (red and orange) are combined into a single group.

**Fig. S7.**
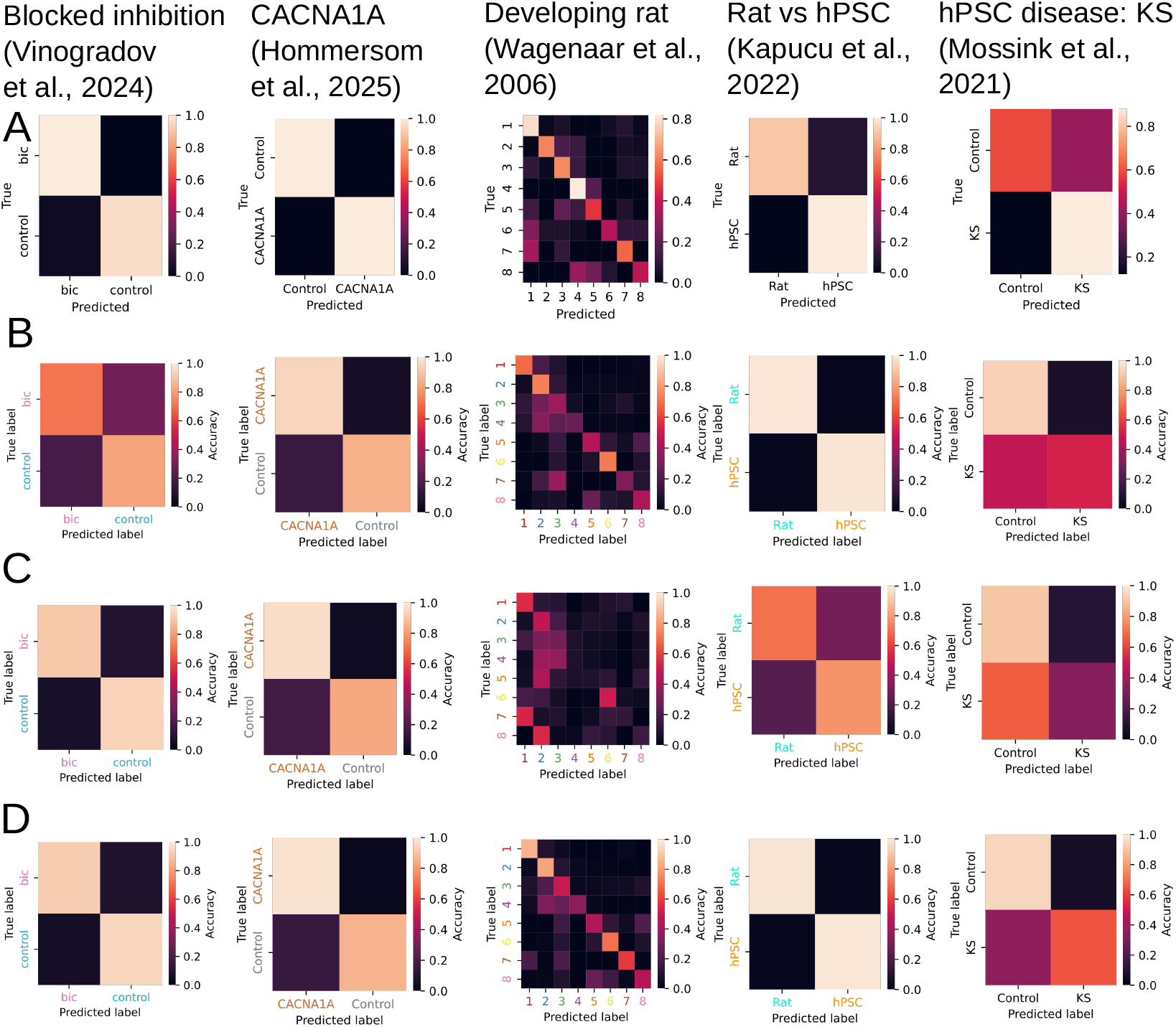
Confusion matrices. **Datasets (columns)**: Mossink et al. [9], Wagenaar et al. [15], Hommersom et al. [25], Vinogradov et al. [26], Kapucu et al. [39] **Algorithms (rows): (A)** KNN Clustering **(B)** XGBoost with traditional burst features **(C)** with burst shape features **(D)** with combined features

**Fig. S8.**
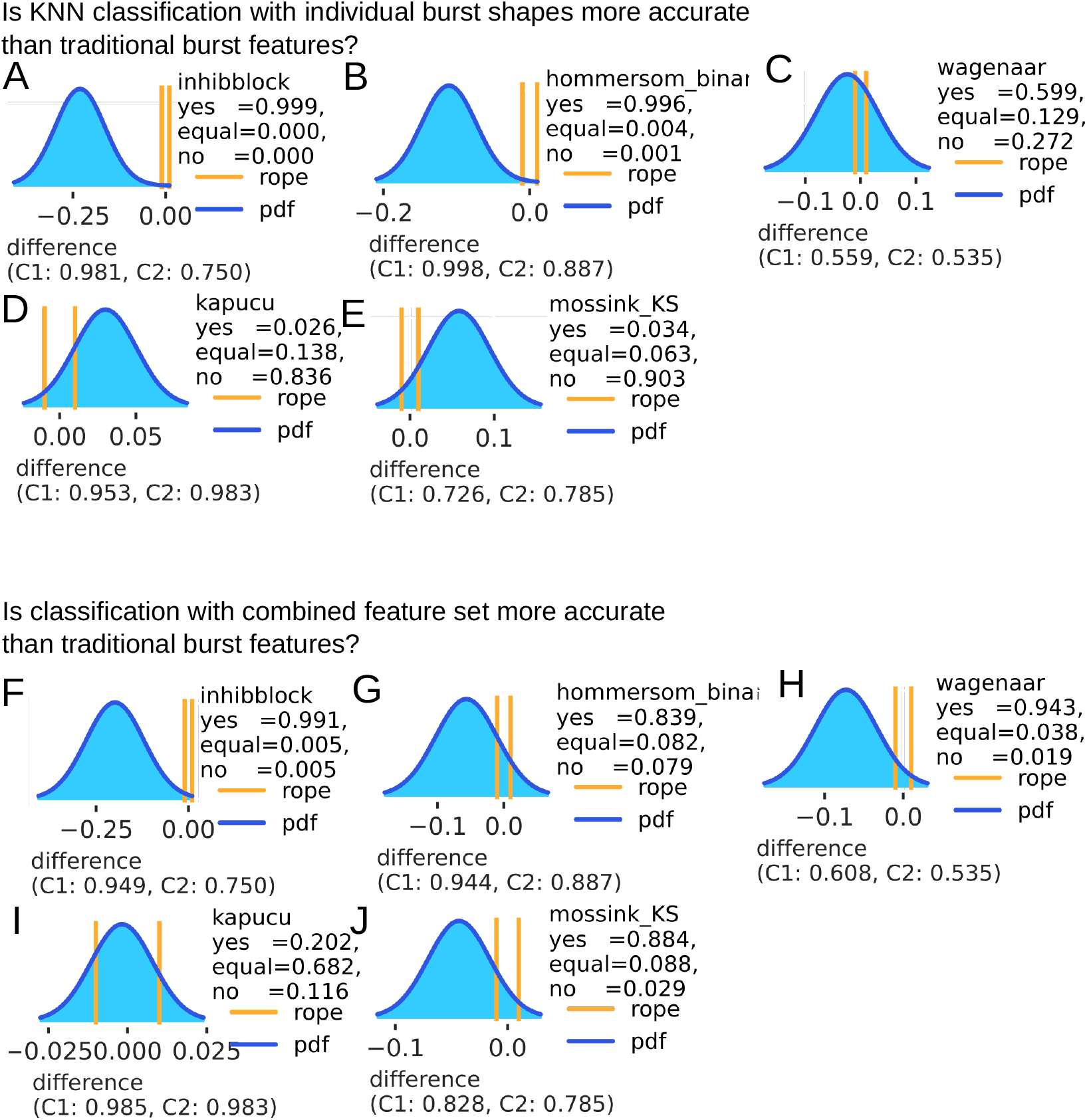
Bayesian comparison of classification accuracies. Bayesian comparison using the baycomp toolbox (https://github.com/janezd/baycomp), with a region of practical equivalence (rope) defined as ±1 % accuracy. The output is the posterior probability distribution that one method is more accurate than the other (see above the *yes* and *no* probability). **(A-E** comparing KNN classification on individual burst shapes against traditional burst shape features. **(F-J)** comparing combined features (shape + traditional) set against traditional burst features.

**Fig. S9.**
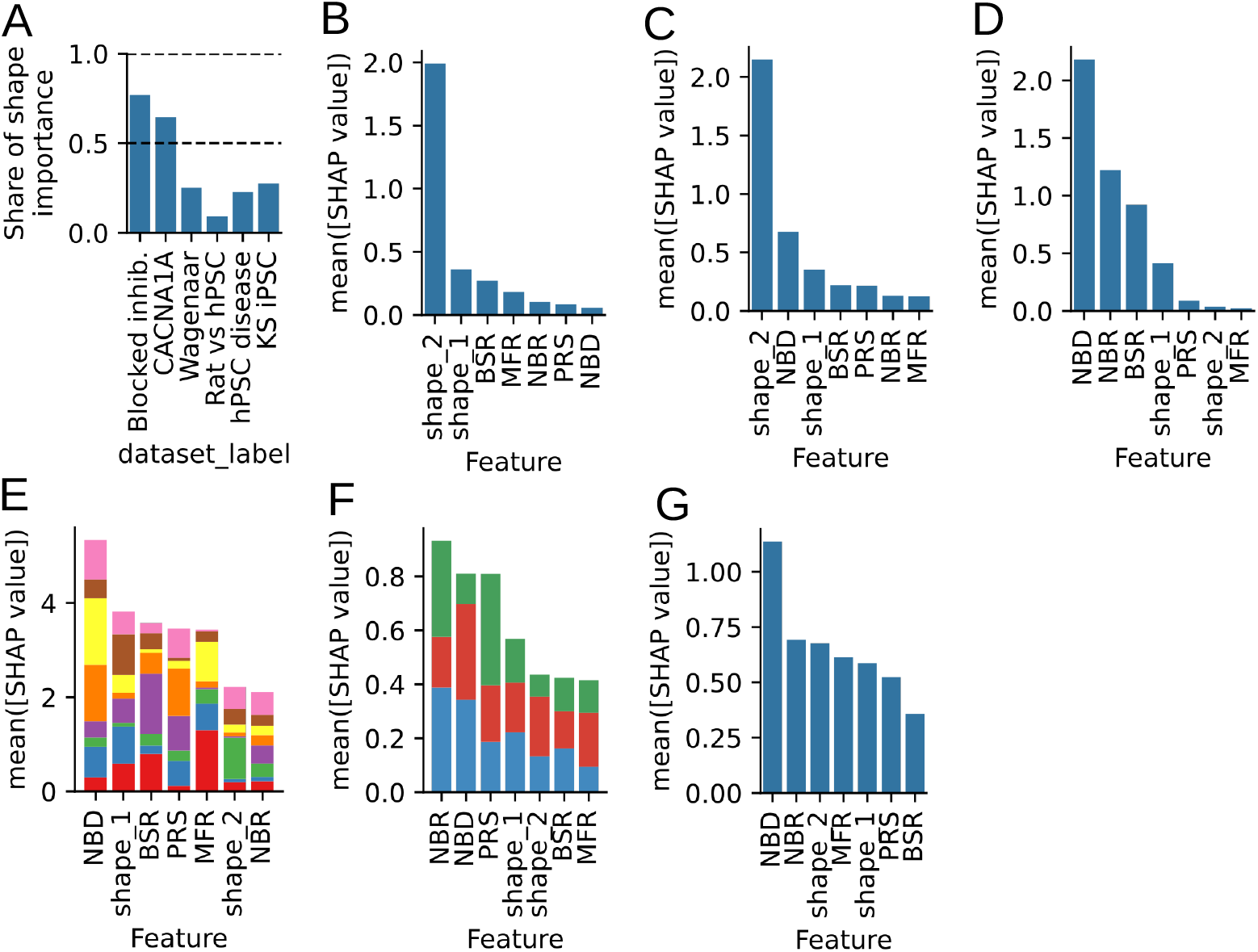
Feature importance measured with Shapley values. **(A)** Summed up feature importance from both shape features (Shapley values are additive). **(B-G)** Feature importance from fitting XGBoost with combined feature set. Order indicates order of importance if more than two groups are present. Colors are group colors. Datasets: **(B)** Blocked-inhibition[26] **(C)** CACNA1A[25] **(D)** Rat vs. hPSC[39] **(E)** Developing rat primary[15] **(F)** Mossink: Kleefstra, MELAS, and Control[9] **(G)** Mossink: only Kleefstra and Control[9].

**Fig. S10.**
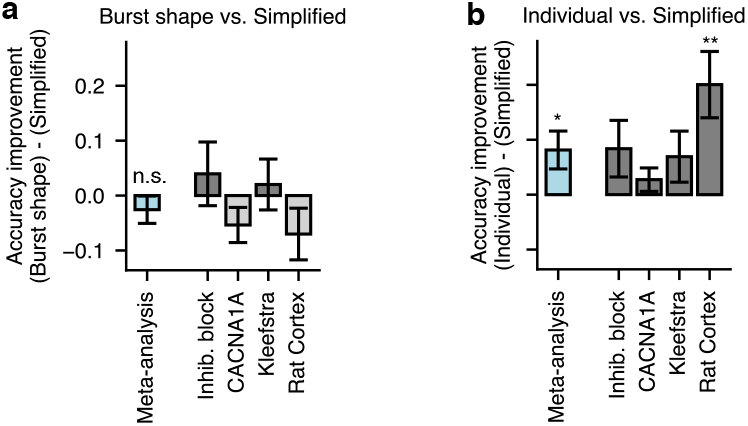
Simplified features (onset and offset) are almost as good as other shape-based classifiers. **(a)** The simplified features perform slightly better than the spectral embedding dimensions (not significant). Note, however, that spectral embedding dimensions can be easily scaled to include more dimensions, whereas the simplified features are manually designed and don’t scale. **(b)** Using the full KNN graph is significantly more accurate than the simplified features.

